# Animal lifestyle changes acceptable mass limits for attached tags

**DOI:** 10.1101/2021.04.27.441641

**Authors:** Rory P Wilson, Kayleigh A Rose, Richard Gunner, Mark D. Holton, Nikki J Marks, Nigel C Bennett, Stephen H. Bell, Joshua P Twining, Jamie Hesketh, Carlos M. Duarte, Neil Bezodis, Milos Jezek, Michael Painter, Vaclav Silovsky, Margaret C. Crofoot, Roi Harel, John P. Y. Arnould, Blake M. Allan, Desley A. Whisson, Abdulaziz Alagaili, D. Michael Scantlebury

## Abstract

1. Animal-attached devices have transformed our understanding of vertebrate ecology. To minimize tag-related harm for these studies, researchers have long advocated that tag masses should not exceed 3% of the animal’s body mass. However, this proposition ignores tag forces generated as a result of animal movement.
2. Using data from collar-attached accelerometers on diverse free-ranging terrestrial animals, we detail a tag-based acceleration method (TbAM) in which we quantify animal athleticism in terms of fractions of animal movement time devoted to different collar-recorded accelerations. The varying accelerations are converted to forces imposed on the animals based on the acceleration and tag mass and allow derivation of defined force limits, including those amounting to 3% of the animal’s mass, for specified fractions of any animal’s active time.
3. We demonstrate how species athleticism is the principal determinant of tag forces, whereas body mass is of little importance. Forces exerted by ‘3%’ tags were mostly equivalent to 4-19% of the animals’ masses during moving, with a maximum of 54% in a hunting cheetah. Cumulative frequency curves of tag acceleration for periods when animals were active, all showed a characteristic sigmoid pattern, which was displaced further to the right as higher acceleration activities accounted for an increasing proportion of any animal’s time. Specifying that tags should exert forces that are less than 3% of the animal’s body mass for 95% of the time led to corrected tag masses constituting between 1.6% and 2.98% of our study animals’ masses, with values depending on animal athleticism.
4. Recognition that animal athleticism affects tag forces of their carriers fundamentally changes how acceptable tag mass limits should be determined by ethics bodies. In order to have a scientifically robust acceptable threshold to limit the forces experienced by an animal carrier, we suggest practitioners derive a similar cumulative acceleration profile for their study species and use a minimum of the 95% limits on the plot (although higher limits may be more appropriate).

## 1. INTRODUCTION

The use of animal-attached devices is transforming our understanding of wild animal ecology and behaviour (Brown et al. 2013; Kays et al. 2015). Indeed, smart tags have been used across scales to measure everything from the extraordinary details of high performance hunts in cheetahs (Wilson et al. 2013), to vast cross-taxon comparisons of animal behaviour and space-use over whole oceans (e.g. (Block et al. 2011; Brown et al. 2013)). A critical proviso is, however, that such devices do not change the behaviour of their carriers, for both animal welfare issues as well as for scientific rigor (Wilson & McMahon 2006). Defining acceptable device loads for animals is critical because even diminishingly small tags can cause detriment. For example, Saraux et al. (2011) showed that the addition of flipper rings to penguins can affect their populations, presumed to be due to the tags increasing the drag force in these fast-swimming birds. Performance is relevant in this case because drag-dependent energy expenditure to swim increases with the cube of the speed (Culik et al. 1993).

Although consideration of the physics of drag has been shown to be a powerful framework with which to understand tag detriment in aquatic animals (e.g. (Rosen et al. 2017; Kay et al. 2019)), drag is negligible in terrestrial (though not aerial) systems even though tag detriment in terrestrial animals has been widely reported and is multi-facetted (Murray & Fuller 2000). Reported issues range from minor behavioural changes (Stabach et al. 2020) through skin-, subcutaneous- and muscle damage with ulceration (Krausman et al. 2004; Hopkins & Milton 2016) to reduced movement speed (Brooks et al. 2010) and dramatically increased mortality (Rasiulis et al. 2014). As with drag, we advocate that a force-based framework is necessary to help understand such detriment. Indeed, force is implicit in ethics-based recommendations for acceptable tag loads because, for example, a central tenet is that animal tags should never exceed 3% or 5% of the animal-carrier body mass (Kenward 2000). Importantly though, we could find no reason for advocating this value, which thus remains arbitrary. Implicit in this limit is that consequences, most particularly the physical forces experienced by animals due to tags, are similarly limited. This cannot be true because Newton showed that mass, force and acceleration are linked *via* F = ma, so animal performance, specifically their acceleration, will affect the tag forces applied to the carriers. Tag forces on the animal carrier can therefore be accessed by measuring acceleration experienced by the tag as the animal moves. We note here though, that this necessitates gathering on-animal data because simple consideration of acceleration from rigid-non-living bodies is inappropriate for living systems composed of multiple interacting segments (Gleiss et al. 2011).

Here, we examine the forces exerted by collar-mounted tags on moving animals. We investigate 4 species within the order Carnivora in detail; lions *Panthera leo*, European badgers *Meles meles*, pine martens *Martes martes* and a cheetah *Acinonyx jubatus* (with body masses roughly spanning 2–200 kg) equipped with accelerometers undertaking their normal activities in the wild for 1-21 days. To help understand how travel speed affects forces, we also equipped twelve domestic dogs *Canis familiaris* (2-45 kg) with the same tags, but with masses of up to 3% of the dog body mass as they travelled at defined speeds. Finally, we equipped a further 6 species of mammal from diverse animal families with different lifestyles with accelerometers *in situ* for 7-168 days. These were: a cercopithecid, the olive baboon *Papio anubis*; a phascolarctid, the koala *Phascolarctos cinereus*; a phalagerid, the mountain brushtail possum *Trichosurus cunninghami*; a bovid, the Arabian oryx *Oryx leucoryx*; a cervid, the red deer *Cervus elaphus* and a suid, the wild boar *Sus scrofa.*

We document how the forces imposed by the collars changed with activity across all these species and conditions. Based on this, we propose a method based on acceleration data that allows researchers to define the breadth of forces exerted by tags on animals and their relative frequency of occurrence. We show how this information can then be used to derive appropriately force-based acceptable limits for tag masses, recognizing the effect of animal lifestyle and athleticism.

## 2. MATERIALS AND METHODS

### 2.1 Tag deployments on free-ranging species

We selected 4 species of free-living carnivores for detailed analysis, exemplifying about 2 orders of magnitude of mass; 10 lions (mean mass *ca.* 180 kg), 1 cheetah (mass *ca.* 40 kg), 10 badgers (mean mass *ca.* 8 kg) and 5 pine martens (mean mass 1.9 kg), and fitted them with collar-mounted tri-axial accelerometers (‘Daily Diaries - Wildbyte Technologies [http://www.wildbytetechnologies.com/]; measurement range 0-32 *g*, recording frequency 40 Hz). Due to the weighting of the loggers, and more particularly their associated batteries, the units and sensors were normally positioned on, or close to, the underside of the collar although during movement the collars could rotate. After being equipped, the animals roamed freely, behaving normally, for periods ranging between 3 and 21 days before the devices were recovered.

In addition to these, we also deployed collar-mounted accelerometers on six select free-ranging animal species to obtain acceleration data from diverse mammal families with varying lifestyles for comparison with the carnivores. These were: a cercopithid, the olive baboon *Papio anubis*; a phascolarctid, the koala *Phascolarctos cinereus*; a phalagerid, the mountain brushtail possum *Trichosurus cunninghami*; a bovid, the Arabian oryx *Oryx leucoryx*; a cervid, the red deer *Cervus elaphus* and a suid, the wild boar *Sus scrofa.* Extensive details on species-specific tagging procedures are included in the Supplementary Materials.

### 2.2 Trials with domestic dogs

Twelve domestic dogs (*Canis lupus domesticus*) of seven different breed combinations and three main body types (small, racers and northern breeds), ranging 2-45 kg in body mass (Table S5), were volunteered by their owners and the RSPCA’s Llys Nini Wildlife Centre (Penllergaer, Wales) to take part in this study (Table S5). Dog body masses were provided by owners and body length, forelimb length and hindlimb length were measured to the nearest cm. Two leather dog collars (short and long) of the same width were used to cover the range in dog neck size. Combinations of pre-prepared lead plates (up to 10 cm in length) and varying in mass (25, 35, 45, 50, 100, 150 and 175 g) were fashioned into collar loads equivalent to 1, 2 and 3% of each carrier dog’s body mass. The loads were stacked and attached securely to the ventral collar along their full-length using Tesa® tape. A tri-axial accelerometer (Daily diary, measurement range 0-32 *g*, recording frequency 80 Hz, Wildbyte Technologies) and its supporting battery (3.2 V lithium ion) were taped securely to the load. The tag and battery combined weighed 28 g and, in the absence of any additional load, were considered negligible in mass and used as a control (0 % carrier body mass). All trials were approved by the Swansea University Animal Welfare Ethical Review Body (ethical approval number IP-1617-21D).

All trials were filmed and each dog was encouraged to walk, trot and bound along a 25 m stretch of level, short-cut grass wearing collar tags equivalent to 0, 1, 2 and 3% of their body mass (twelve gait and tag mass combinations) and trial order was randomized. A stopwatch was used to record the time taken (to the nearest s) for a dog to travel 20 m between appropriately spaced posts in order to calculate an average speed of travel (m s^−1^).

### 2.3 Data processing

In both the cases of the free-living carnivores and domestic dogs, the 3 channels of raw acceleration data were converted to a single channel by calculating the vectorial sum of the acceleration following 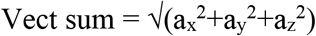, where a is the instantaneous acceleration and the subscripts denote the different (orthogonally placed) acceleration axes. The specifics of the surge, heave and sway accelerations were not considered separately due to some collar roll. We selected 4 peak accelerations from the gait waveforms to examine as a function of speed, gait, body mass and tag mass as a percentage of carrier body mass in the dogs (SI 2). We standardized the use of four peaks because at the highest speeds some dogs only had four full waveforms during the test stretch. Gait was assessed visually in the dogs as a walk, trot or gallop. The forces exerted by the tags on their animal carriers were calculated using F = ma, where m is the mass (kg) of the tag and a is the acceleration (*g*).

### 2.4 Tag-based acceleration method (TbAM)

Finally, in a full cross-species comparison, we took the vectorial sum of the tri-axial acceleration data (see above) and plotted the cumulative frequency distribution from each species to define the vector sum of the acceleration at species-specific 95% and 99% limits.

### 2.5 Statistical analyses

Linear mixed-effects models were conducted in R Studio V 1.2.1135 within the ‘Lme4’ package in order to investigate how the period between acceleration peaks, gait and body mass influenced peak accelerations across species. Additionally, we investigated how travel speed (covariate), body mass (covariate), collar mass as a percentage of carrier body mass (fixed factor) and gait (fixed factor) influence peak accelerations and consequent forces exerted by the tags. The influence of dog body mass and collar mass as a percentage of carrier body mass on gait-specific travel speed and the period between peak accelerations was also investigated. Dog ID was included as a random factor in all models. All potential interaction effects were first investigated and a step-wise back-deletion of non-significant interaction terms was conducted. Standard model diagnostics were conducted in order to ensure that model assumptions were met (examining q-q plots and plotting the residuals against fitted values) and data transformations were conducted in order to meet assumptions where appropriate. Outputs of the final models are reported. The F statistic and marginal and conditional R^2^ were determined using the ‘car (3.0-5)’ and ‘MuMIn (1.46.6)’ packages, respectively. Coefficients for best-fit lines in the figures were extracted from the outputs of the models

## 3. RESULTS

### 3.1 Changing acceleration according to activity in carnivores

Accelerometer data for periods when our carnivores travelled, summarized as the vectorial sum of the three orthogonal axes, showed tri-modal distributions except for the pine martens which were mono-modal. Following (Dewhirst OP et al. 2016) we considered that these most likely corresponded to walking, trotting and bounding (e.g. Fig 1, cf. our direct observations of the domestic dogs below); these were further exemplified by variation in the amplitude and periods of peaks in this acceleration metric (Fig. 2). Cumulative frequencies of all acceleration values showed increasing acceleration from walking through trotting to bounding and typically had a roughly logarithmic-type curve for all gaits and animals (Fig. 1). The percentage time during which the tags carried by the carnivores had acceleration exceeding 1 *g* varied between a mean minimum of 31% for walking badgers to 88% for bounding cheetahs (Table S1). Furthermore, while differences in species acceleration distributions were not readily apparent for their walking gaits, the percentage time during which acceleration was in excess of 1 *g* was greatest during bounding, with cheetahs showing the highest values in this category (Fig. 1). Mean peak accelerations per stride across species varied between 1.37 *g* (SD 0.05) and 6.25 *g* (SD 0.79) for walking and bounding cheetahs, respectively (Table S2). The maximum recorded value was 18.1 *g* in a cheetah assumed to be chasing prey.

**FIGURE 1.**
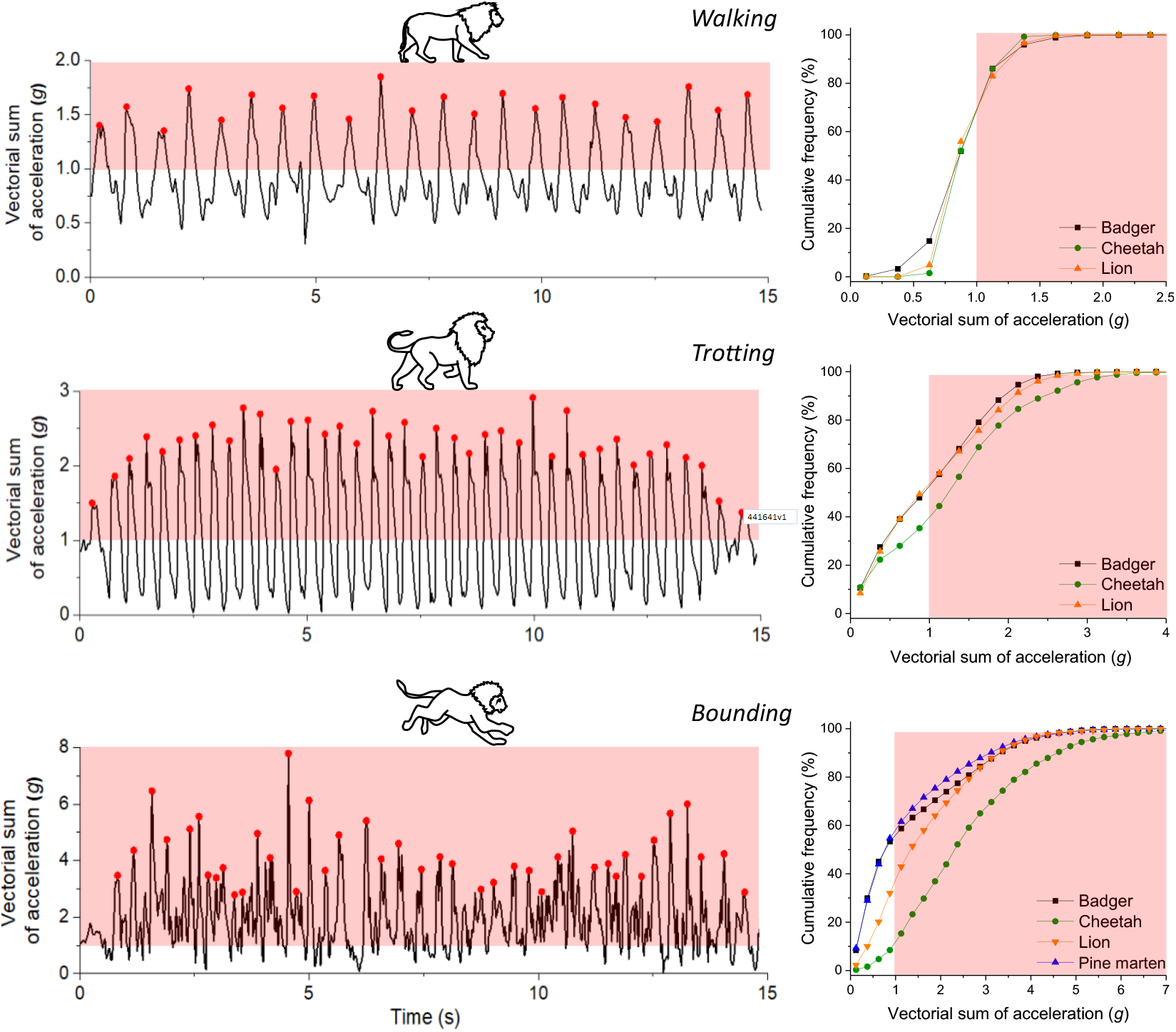
Acceleration signatures vary according to gait and lifestyle. Left-hand panels; Acceleration signatures recorded by collar-mounted tags on a lion according to activity. The red areas show when the acceleration exceeded that of gravity (note the changing scales with gait). Right-hand panels; Cumulative frequency of all acceleration values for four free-living carnivores according to gait. Note that the pine martens never walked or trotted.

**FIGURE 2.**
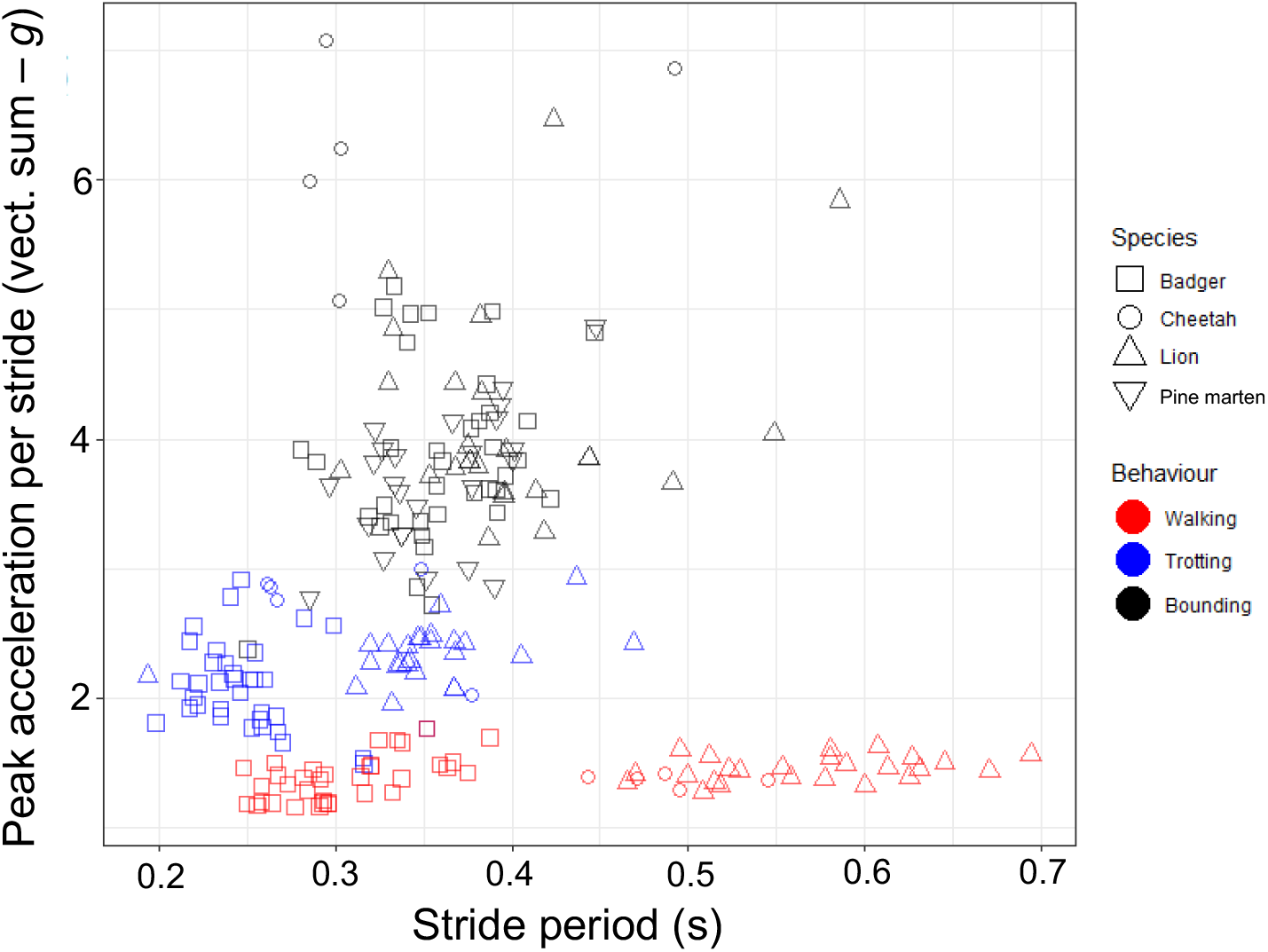
Body mass and stride period do not dictate peak tag acceleration. Distributions of peak amplitudes of (the vectorial sum of) accelerations and stride periods for four free-living carnivores (see symbols, with mean masses of *ca.* 2 kg, 8 kg, 40 kg and 180 kg for the pine martens, badgers, cheetah and lions, respectively) travelling using different assumed gaits (colors). Each individual point shows a mean from a duration of activity >5 s from a single individual. See also Fig. S1 for similar data from domestic dogs.

Across species, gait was the main factor dictating peak acceleration (Fig. 2) and there were no significant effects of body mass, nor period between peaks (linear mixed-effects model: log period: *F*_1, 210_= 0.01, *P*=0.908; gait: *F*_2, 208_=1083.07, *P*<0.0001; body mass: *F*_1, 19_= 3.00; *P*=0.100, Table S3). The period between acceleration peaks was greater for larger species during slower gaits, but not for bounding (a linear mixed-effects model demonstrated a significant interaction effect between body mass and gait: *F*_2, 209_= 3.00, *P*<0.0001, Table S3).

Acceleration metrics changed even within particular gaits though, as exemplified by predators chasing prey. For example, overall, prey chases exhibited by lions showed mean peak accelerations increasing from 1 *g* to mid-chase peaks of *ca.* 5 *g* before decreasing again (Fig. 3). Assuming that these animals were carrying tags that amounted to 3% of their mass, this would result in forces amounting up to and beyond 15% of their normal body mass (e.g. Fig. 3). The maximum observed was 54% in the cheetah.

**FIGURE 3.**
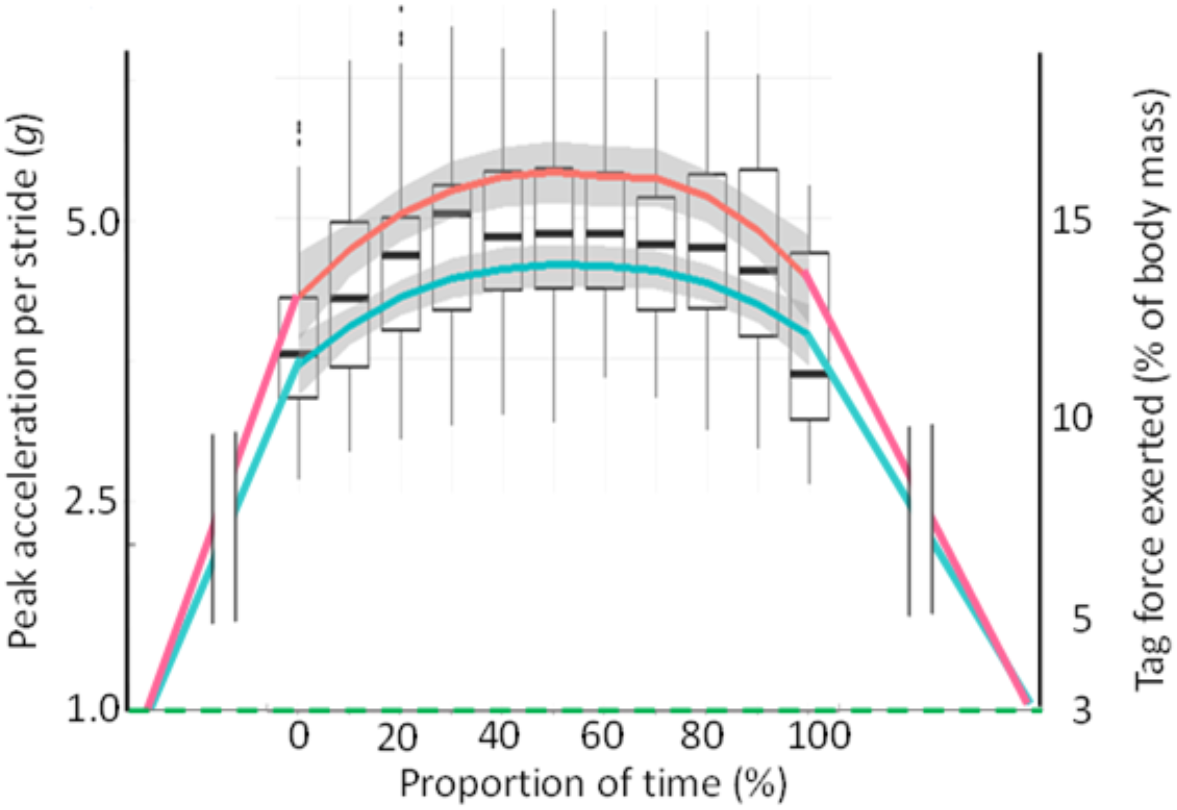
Hunting lions experience maximum tag forces mid-chase. Box and whisker plots [bold horizontal bars show means, boxes inter-quartile ranges and whiskers 1.5 X IQR] of the changing forces (the vectorial sum of the acceleration peaks per bound [cf. Fig. 1] and as a percentage of animal body mass assuming the tag constitutes 3% of body weight during non-movement) exerted by animal-attached tags on lions chasing prey as a function of the progression of the chase. Red and blue lines show grand means for 5 females and 5 males, respectively. The maximum acceleration was > 12.5 g, equating to a 3% tag exerting a force equivalent to 37.5% of the animal’s normal weight.

In dogs, stride peak accelerations increased linearly with travel speed (Fig. 4), but at greater rates with increasing relative tag mass above tag masses equivalent to 1% of carrier body mass (there was a significant interaction effect between travel speed and tag % body mass: *F*_3, 500_ = 4.44, *P*=0.004, *R*^2^ = 0.74 Table S4). There was also a significant interaction effect between gait and tag mass as a percentage of carrier body mass (*F*_6, 498_ = 4.33, *P*=0.0002, *R*^2^ = 0.74, Table S4). Peak accelerations ranged from 4-18 *g* (Fig S1-2) during bounding with collar tags equivalent to 3% of the carrier body mass. Movement of the tag relative to the collar and body (flapping/ swinging) was exacerbated under this condition and, as a consequence, the force exerted by the tags ranged from 20-50% of normal carrier body mass.

**FIGURE 4.**
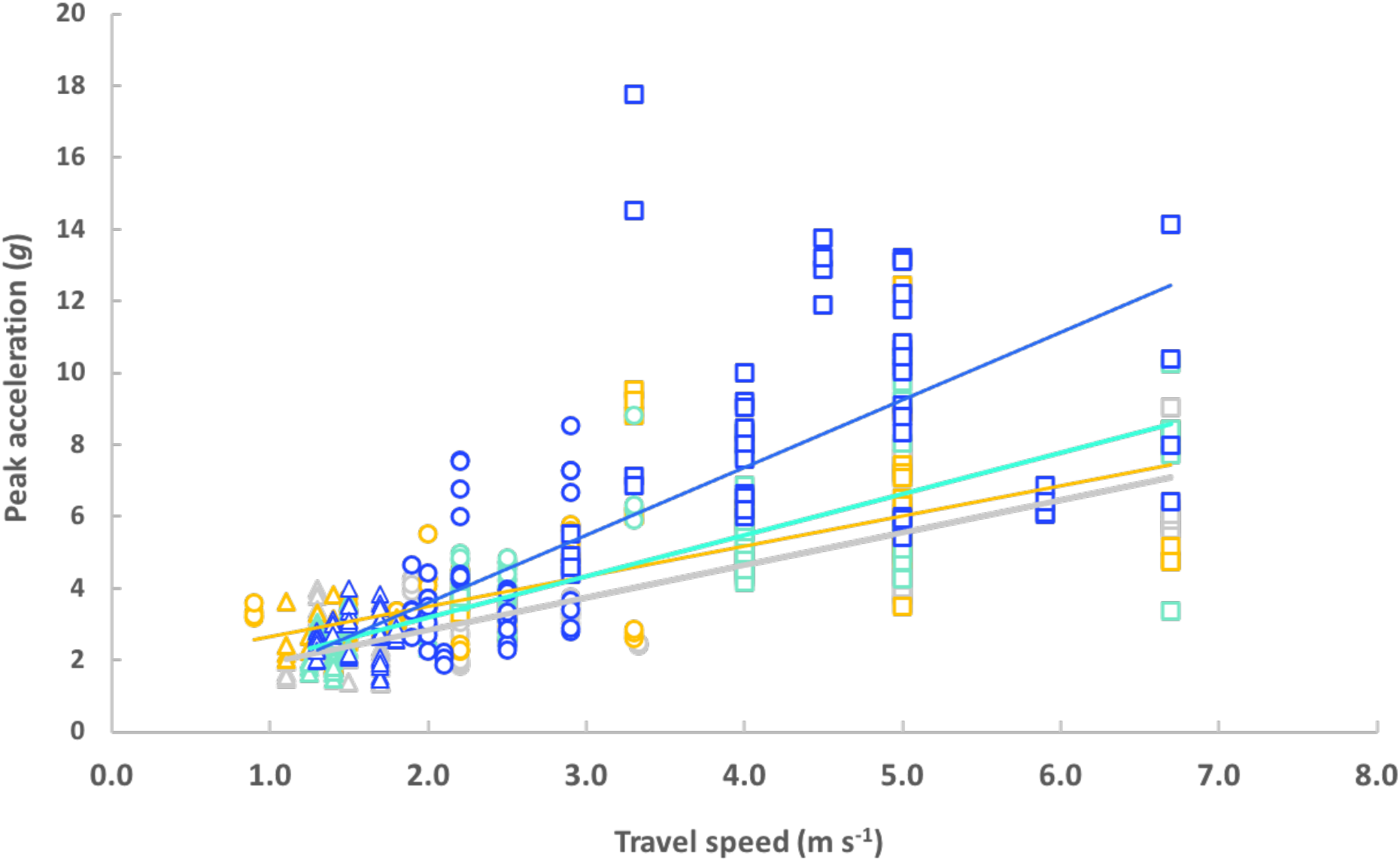
Travel speed and tag mass influence tag peak acceleration. The relationship between peak accelerations and travel speed for 12 individual dogs (masses 2-45 kg), colored according to the percentage mass of the tag relative to the carrier; 0% (grey, *y = 0.90x + 1.04*), 1% (yellow, *y = 0.84x +1.80*), 2% (light blue, *y = 1.15x + 0.88*), 3% (dark blue, y = 1.88x - 0.16). Data points are the means of the four greatest peaks in acceleration per 20 m trial per gait and dog. Data for walking, trotting and bounding gaits are represented as triangles, circles and squares, respectively. Coefficients for the best-fit lines are taken from the final model outputs (Table S4).

Stride peak accelerations were largely invariant with body mass (*F*_1,10_ = 3.51, *P*=0.09, Table S4) across dog breeds for any given gait (Fig. S2). Consequently, the peak forces exerted by the tags were directly proportional to tag mass and body mass. Accordingly, relative forces (force as a percentage of normal carrier body mass) were independent of carrier body mass (Table S4, Fig. S3).

### 3.2 Using accelerometry to derive an over-arching tag-force rule

Although travelling is a major component across species, animal activity across all behaviours contributes to the acceleration, and therefore the tag force profiles, that animals experience. We produced cumulative frequency curves of the vectorial sum of the acceleration (cf. Fig. 1) for all 10 study species for periods when animals were considered active and these all showed a characteristic sigmoid pattern (Fig. 5a). These relationships were displaced further to the right as higher acceleration activities accounted for an increasing proportion of any animal’s time (Fig. 5a). In order to have a scientifically robust acceptable threshold to limit the forces produced by a tag on an animal carrier, we suggest a tag-based acceleration method (TbAM); that researchers should derive a similar cumulative acceleration profile for their study species and use a minimum of the 95% limits on the plot (although higher limits may be more appropriate). Assuming, in the case of our study animals, that these limits were intended to cater for a tag that should exert forces that are less than 3% of the animal’s body mass, this limit would lead to corrected tag masses constituting between 1.6% and 2.98% of our study animals’ masses (Fig. 5b). We note however, that even these corrected tag masses would effectively exceed the 3% rule conditions for 1/20^th^ of the animals’ active periods: The difference between the 95% and 99% thresholds for our study species indicates the extent of the force development for this period with some, such as the koalas, showing virtually no difference, whereas badgers, baboons and martens exhibited substantial differences (Fig. 5b). Importantly though, this method would allow researchers to define any tag force thresholds, not just 3%, and the times these were exceeded by the animal, not just 95%.

**FIGURE 5.**
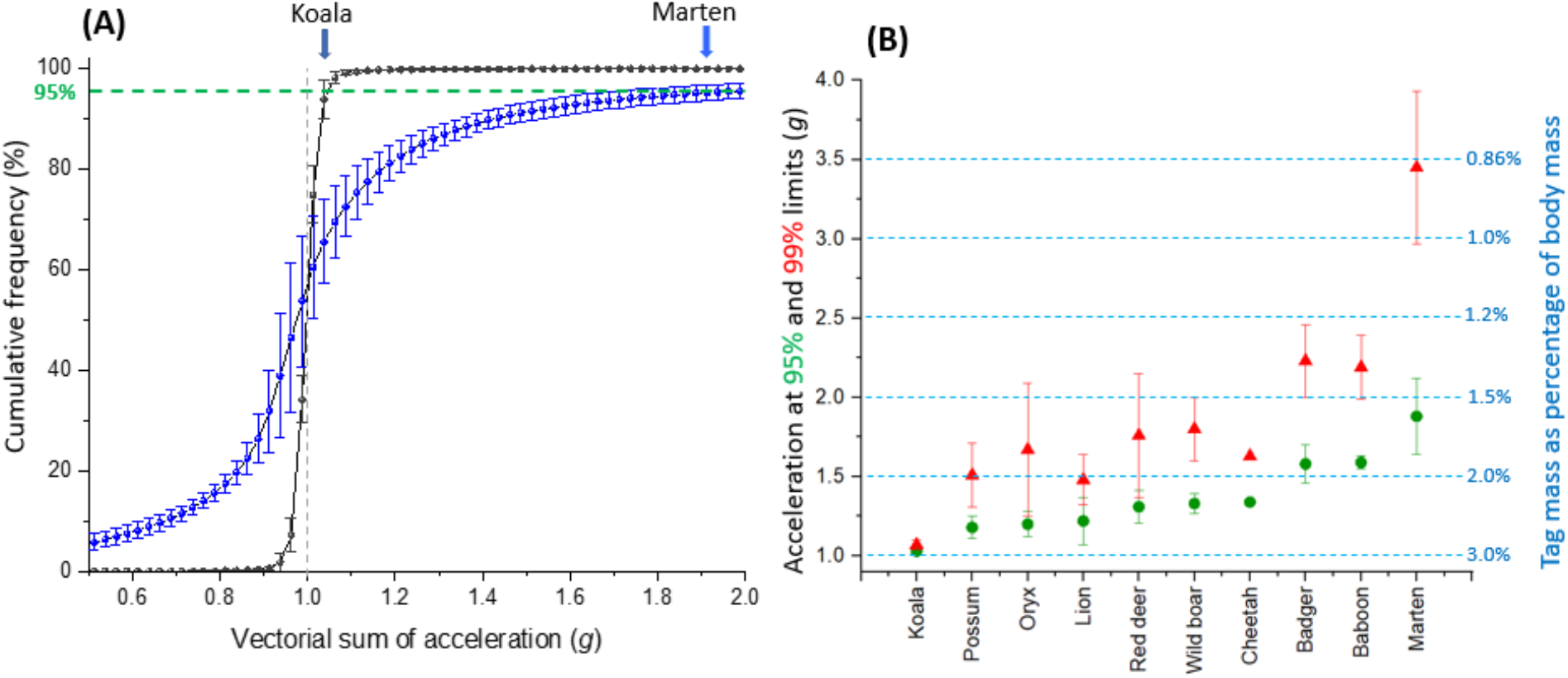
Defining tag mass limits based on cumulative time spent experiencing tag forces. (A) shows the mean cumulative frequency (bars = SD) of the vectorial sum of the acceleration for two arboreal animals with very different athleticisms: 5 koalas (black line) and 5 pine martens (blue line). The 95% limit is shown by the dashed green line and the relevant points are shown for both species by colored arrows. (B) shows these two species points (and the 99% limits in red) adjacent to a broader species list highlighting variation in lifestyles. Assuming that a tag should only exert a force amounting to 3% of the carrier animal’s mass, the translation of these species-specific acceleration limits can be used to correct tag masses to be an appropriate percentage of the carrier animal mass (blue axis on the right).

## 4. DISCUSSION

A rigid vehicle accelerating in a straight-line only experiences acceleration in the longitudinal axis. In contrast, the multiple limb-propelled motion of an animal with a flexible body produces complex three-dimensional trunk accelerations owing to the changing limb accelerations (Gleiss et al. 2011) caused by multiple muscle groups that ultimately transfer mechanical energy and affect shock absorption (Wu et al. 2019), and the mechanical work conducted within each stride (Biewener 2006). Ultimately, the magnitude of trunk accelerations depends on the combined acceleration of the limbs, and the masses of those limbs (cf (Gleiss et al. 2011)). Thus, animals engaging in high performance activities are expected to produce high body accelerations, and have physiological and anatomical adaptations to enhance performance, such as fast twitch muscles (Gregor et al. 1979), and tendons designed for greater storage and release (Alexander 2002), which will increase this. Through all these complexities, tags mounted on the trunk of an animal result in greater forces being imposed that scale linearly with the acceleration of the tag and its mass. Consideration of animal lifestyle then, can already inform prospective tag users of the likely scale-up of the tag forces beyond the 1 *g* normally considered for tag detriment. Consequently, the 3-5% mass limits for slow-moving animals, such as sloths (Bradipodidae) or koalas (Phascolarctidae) (Fig. 5), seem most appropriate, while they may not be for pursuit predators, such as wild dogs (*Lycaon pictus*), regularly jumping animals like kangaroos (Macropodidae) or martens (Mustelidae) (Fig. 5) and rutting ungulates (Ungulata). Beyond that, in our small sample of carnivores at least, which nonetheless covers about two orders of magnitude in mass, it seems that peak acceleration associated with gait varies little with mass, although larger animals have longer stride periods (Fig. 2 – cf. (Farley et al. 1993)). If these animals were to carry tags constituting 3% of their normal body mass, mean peak forces imposed by the tags would constitute *ca*. 4.5%, 6% and 12% of this body mass for walking, trotting and bounding gaits at frequencies of between 1.6 and 4 times per second (for walking lions and trotting badgers, respectively - Fig. 2, Table S2).

Importantly, tag attachment is relevant in translating the acceleration experienced by the animal’s trunk into tag-dependent forces acting on the animal, with collars predicted to be particularly problematic. A tag that couples tightly with its carrier’s trunk, such as one attached with tape to a bird (Wilson et al. 1997) or glue to a marine mammal (Field et al. 2012), experiences acceleration that closely matches that of its substrate, so it exerts forces at a site where most of the animal’s mass lies. In contrast, a device on a looser-fitting collar of a moving tetrapod not only exerts forces on the (less massive) head and neck areas, rather than the animal trunk, but the tag also oscillates between essentially two states: One is analogous to ‘freefall’, which occurs between pulses of animal trunk acceleration in the stride cycle which project the collar in a particular direction owing to its inertia and lack of a tight couple with the neck. The collar is therefore subject to peaks in acceleration when it interacts with the animal’s neck, causing greater collar acceleration than would be the case if it were tightly attached to the animal’s body (cf. peaks in Fig. 1). This explains why Dickinson et al. (Dickinson et al. 2020) reported that acceleration signatures from collar-mounted tags deployed on (speed-controlled) goats *Capra aegagrus* became increasingly variable with increasing collar looseness, and is analogous to the concerns related to injuries sustained by people in vehicles depending on seatbelt tightness (Hodson-Walker 1970). Partial answers to minimizing such problems may involve having padded collars that should reduce acceleration peaks, making sure that the tags themselves project minimally beyond the outer surface of the collar and having wider collars to reduce the pressure.

Having identified how animal movement changes the 3% tag rule, it is more problematic to understand how the identified forces translate into detriment. A prime effect is that higher forces and smaller contact areas will lead to higher pressure at the tag-animal interface because pressure = force/area. This can affect anything from fur/feather wear (Young & Blob 2016) to changing the underlying tissue (Michael et al. 2013) and, as would be predicted, is notably prominent in species wearing thin collars (e.g. Howler monkeys *Alouatta palliatai*, where 31% of animals wearing ball-chain radio-collars constituting just 1.2% of their mass sustained severe damage extending into the subcutaneous neck tissue and muscle (Hopkins & Milton 2016)). But pressure-dependent detriment will also depend on the proportion and length of time to which an animal is exposed to excessive forces, with animals that spend large proportions of their time travelling, such as wild dogs, being particularly susceptible (Pomilia et al. 2015).

Perhaps more esoteric, is the extent to which the inertia of a variable force-exerting tag ‘distracts’ its wearer, aside from the physical issues of load-bearing by animals. The tag mass as a percentage of carrier mass did not affect the gait-specific speeds selected by the domestic dogs in this study. However, it remains to be seen the extent to which a typical 30 kg cheetah wearing a collar that is 3% of its body mass, and therefore experiencing an additional force equivalent to up to 16 kg during every bound of a prey pursuit, might have its hunting capacity compromised. We note that the survival of such animals is believed to be especially sensitive to the proportion of successful hunts (cf. (Scantlebury et al. 2014)) which calls for critical evaluation of performance between tag-wearing and unequipped animals, or animals equipped with tags of different masses (cf.(Wilson et al. 1986)).

In the meantime, our suggested approach of setting tag mass limits based on the overall (corrected) forces being less than 3% of the animal’s mass for 95% of the time should go some way to getting a more realistic assessment of the potential for detriment. Where researchers adopting this approach do not have appropriate acceleration data for their study animal, they could use a surrogate species, perhaps from an online database. Such a resource should define the length of time that study animals were equipped to derive the acceleration frequency distribution as well as note other pertinent factors such as season, that might affect the distribution.

Importantly, we do not advocate the 3% rule as such, but recognize that it has been widely adopted and could serve as a useful starting point with which to consider tag detriment if calculated as we have suggested here. In this, cognizance should also be given to the extent of tag forces for periods above the 95% threshold because, where these are excessive, it may be appropriate to use a 99% threshold or higher to derive appropriate tag masses. Notably though, even 99% limits do not highlight the high tag forces developed during prey pursuits exhibited by the cheetah. We suggest that the solution to this lies in more detailed consideration of the animal’s lifestyle, in particular identifying survival-critical behaviours with exceptionally high accelerations. Such periods may persuade ethics bodies to raise their thresholds still further. Underpinning this will be ongoing miniaturization, where tags benefit from the sensor revolution in human wearables, which will undoubtedly percolate through to animal applications: Advanced smart phones have >10 sensors, along with significant memory, battery and data transmission capabilities, and typically weigh 150 - 200 g or about 0.2 % of average human body mass. There is, therefore, no reason why scientists and supply companies should continue to use a 3% body mass as a reference *per se* (cf (Portugal & White 2018)).

Finally, consideration of the acceleration-based forces generated by animal-attached tags does not cover all forms of detriment because other forces are at play, such as greater drag in swimming- and flying species (cf.(Saraux et al. 2011)), and more esoteric elements, such as device colour, that affect animal behaviour (Wilson et al. 1990). However, our framework should take the current ‘one-size-fits-all’ basic 3% rule into an arena where quantitative assessment of acceleration can be compared to the myriad of tag-influenced behaviours recognized by the community to link animal lifestyle to putative detriment. Most importantly, these considerations should give ethics bodies a more useful rule of thumb than is currently the case and enable us to develop systems that minimize force-based tag effects, to the benefit of both animals and the science that their studies underpin.

## ACKNOWLEDGEMENTS

We gratefully acknowledge the access provided by the National Trust and Forest Service NI. We are also grateful to the RSPCA’s Llys Nini Wildlife Centre in Penllergaer, Wales, and to Judy Corbett and Peter Welford at Gwydir Castle in Llanrwst, Wales, for allowing us to work on their property with their dogs. We thank Derek van Heerden and the staff at Harnas Wildlife Foundation in Namibia for their kindness and supporting our work. We thank SANParks and the Department of Wildlife and National Parks, Botswana for allowing our research in the Kgalagadi Transfrontier Park (Permit Number SCAM 1550). We are grateful to Angela Bruns, Sam Ferreira and Danny Govender and Pauli Viljoun for facilitating the research and to the many field staff and volunteers that conducted the fieldwork including Wayne Oppel, Corera Links, Martin van Rooyen and Mads Frost Bertelsen. We are grateful to Fraser Menzies and all the field staff at the Department of Agriculture, Environment and Rural Affairs, DARD. We are also grateful to the Vincent wildlife trust for supporting this research. AA, DMS and NCB are particularly grateful to Prince Bander bin Saud Al-Saud, past President of the Saudi Wildlife Authority (SWA), for his unlimited and enthusiastic support to undertake these studies on the Arabian oryx, managed by the SWA

## AUTHORS’ CONTRIBUTIONS

Conceptualization: RPW, KARR

Methodology: RPW, KARR

Investigation: RG, SHB, NM, JPT, JH, JA, BMA, DAW, NCB, MCC, RH, VS, MP, MJ, AA, DMS

Software development: MH

Visualization: RPW, RG, MH, KARR

Supervision: RPW, KARR

Writing—original draft: RPW, KARR, DMS

Writing—review & editing: All authors

## DATA AVAILABILITY STATEMENT

All data will be made available on figshare

## SUPPORTING INFORMATION

Additional supporting information may be found online in the Supporting Information section

