## Supplementary information for "Animal lifestyle changes acceptable mass limits for attached tags"

**Supplementary Materials**

Details on species-specific tagging procedures

Badgers were live-trapped in various locations in Northern Ireland in custom-built cages and anaesthetised with a mixture of ketamine, medetomidine and butorphanol IM with recumbency achieved 5-10 minutes post injection. Loggers were attached to an adjustable nylon clip-on dog collar (Ancol Pet Products Limited, Walsall; circumference 20-30 cm) with a layer of waterproof self-amalgamating tape (ultratape; Bruce Douglas Marketing, Dundee, UK), which was additionally fastened with three cable ties and then covered with 'tesa' tape (No. 4651; tesa AG, Hamburg, Germany). Badgers were recaptured approximately 10 days later, and the collars removed.

Live trapping and collar deployment was conducted at two sites under UK Home Office License and Northern Ireland Environment Agency, License 2228 between August and November 2019. Pine martens were lived trapped in the Crom Estate and Slieve Gullion in Northern Ireland using Tomahawk 205 live cage-traps (Tomahawk Live Trap Ltd., USA). Trapped animals were anaesthetised with an intramuscular injection of ketamine (25 mg per kg) and midazolam (0.2 mg per kg body mass). Loggers were made waterproof and shockproof using Guronic casting resin (C500, SE Connectivity, Schaffhausen, Switzerland) and were deployed on custom-made chrome leather collars (circumference, 11 – 22cm) with self-amalgamating tape (RS PRO Black Self Amalgamating Tape 25mm x 10m, RS components, Northants, UK). The complete system amounted to a collar weight of approximately 45 g (*ca.* 2.3% of bodyweight). Deployments lengths ranged from 5 to 14 days.

The cheetah work took place at Harnas Wildlife Foundation in Namibia. A hand-reared individual was used, so a collar could be placed around its neck without anaesthesia. Once released, the individual was free to move and hunt at will. Data were downloaded each day for a total tagged period of 3 days. We used a commercially available nylon collar (EzyDog Medium dog collar, circumference 29-40cm, Pets at home, Handforth, England) to which we attached the logger with three cable ties and tesa' tape as described above.

Lions were captured from the Kgalagadi transfrontier park in South Africa and treated according to SANParks operational procedures as detailed in SANPark's 'Standard Operating Procedures for the Capture, Transportation and Maintenance in Holding Facilities of Wildlife' (Reference: 17/Pr-CSD/SOP capture, transport, holding facilities (04-17) v2) and SANParks SOP 'Fitment of tracking devices and marking of fauna in South African National Parks' (Reference: 17/Pr-CSD/pro/tracking + marking (12/16) v1). A pride of lions was identified during the course of the day through field rangers' observations and tourist sightings. At nightfall a bait station was set up nearby the pride, consisting of an antelope carcass secured to a tree. This carcass was freshly acquired during the day by authorized ranger staff. The intestines were removed via a cut through the abdominal midline and intestinal contents used to lay a scent trail towards the bait. The darting vehicle was positioned at 20 m distance from the bait station and lion alerted and lured towards the bait by broadcasting pre-recorded animal distress calls via a long-range loudspeaker system (Truck Pro, Foxpro Inc). Once lions settled to feed, the targeted individuals (all adults) were identified and their weight estimated. Subsequently the animals were darted with a DAN-INJECT CO2 injection rifle (Model JM.SP.25, DAN-INJECT). A drug combination of zolazepam/tiletamine (Zoletil®Virbac) and medetomidine hydrochloride (Medetomidine Compound, Kyron Laboratories,) at an average 1.2 mg/kg and 0.05 mg/kg respectively as adapted from recommendations in SANPark's 'Standard Operating Procedures for the Capture, Transportation and Maintenance in Holding Facilities of Wildlife' and Kock *et al. (34)* was used. The darted individuals were followed visually with the help of spotlights. Once animals became recumbent they were recovered, secured with blindfold and foot shackled and translocated to a nearby processing station. They were monitored for temperature, respiration and heart rate. Collars were fitted around the neck in accordance with SANParks SOP 'Fitment of tracking devices and marking of fauna in South African National Parks', allowing for three fingers space and ensuring collar size did not exceed the maximum head circumference. Where prolonged anaesthesia was required, animals were given an additional increments of ketamine (Ketamine Powder Compound, Kyron) intramuscularly at a total average of 1.44mg/kg. Dart sites were treated systemically via subcutaneous injection at the recommended dose with Ceftiofur (Excede®, Zoetis,, 1mL per 30 kg) and meloxicam (Metacam®, Boehringer, Randburg, 0.2mg per kg) respectively. Once processed, animals were relocated close to the pride or capture location and the medetomidine component of anaesthesia reversed intramuscularly with a mix of atipamezole (Antisedan®, Zoetis) at 2.5 times the amount of medetomidine and yohimbine (Yohimbine Compound, Kyron) at 6.25 mg per kg body weight. Animals were monitored until ambulatory. All procedures followed recommendations of SANPark's 'Standard Operating Procedures for the Capture, Transportation and Maintenance in Holding Facilities of Wildlife'. Pine martens (males) were live-trapped and equipped with tags in Northern Ireland between November 2018 - March 2019, and August - November 2019. Devices were attached to the collar using self-amalgamating tape. The complete system amounted toa collar weight of approximately 45g (*ca.* 2.3% of bodyweight). Deployments lengths ranged from 5 – 14 days.

The Arabian Oryx work took place at Mahazat as-Sayd, a large protected area in west-central Saudi Arabia (28°15' N, 41°40'E). Individuals were captured during August 2014 and February 2015. Animals were remotely injected using a Dan-Inject dart gun (Daninject, Børkop, Denmark) with etorphine hydrochloride (CaptivonTM 98, Wildlife Pharmaceuticals Ltd., White River, South Africa; 24µg/kg), Ketamine (Ketaminol® Vet., MDS Animal Health, Intervet International B.V., Boxmeer, The Netherlands; 0.3mg/kg), Midazolam (Midazolam, Wildlife Pharmaceuticals Ltd., White River, South Africa; 0.13mg/kg), and Medetomidine (Zalopine 10 mg/ml, Orion Pharma, Espoo, Finland; 6µg/kg), as described by *(36).* Whilst recumbent, animals were fitted with neck collars to which Daily Diary tags were attached with cable ties and a 200g weight, which ensured that the tag remained at the front and bottom of the collar. After the procedure, anaesthesia was reversed using naltrexone hydrochloride (Naltrexone, APL, Kungens Kurva, Sweden; 40mg IM) and atipamezole hydrochloride (Antisedan, Orion Pharma; 2mg IM) and animals were allowed to recover in an outside shaded enclosure (25 x 25m), before being released. Ten days post deployment, the animals were recaptured and sedated using the methods described above, the collars were removed and the data from the tags were downloaded.

Olive baboon fieldwork was conducted at the Mpala Research Center, a conservancy consisting of nearly 200 $km^2$ of savannah and dry woodland habitats in central Kenya. All research activities described in this paper were approved by the Republic of Kenya's National Council for Science and Technology (Permit # NCTS/RCD/12B/012/26B), Kenya Wildlife Services

(KWS/BRM/5001) and the Smithsonian Tropical Research Institute's Animal Care and Use Committee (IACUC # 2012.0601.2015). From July 21st – 29th, 2012, 33 baboons were captured using two arrays of individual traps (1 $m^3$), which were baited with maize and placed at sites near the troop's sleeping trees. Baboons were chemically immobilized using Ketamine (15 mg/kg). 26 baboons were fitted with GPS collars (e-Obs Digital Telemetry, Gruenwald, German). Adults and large subadults were fitted with D-cell battery collars weighing 300 g equipped with a break-away mechanism (Advanced Telemetry Solutions, Isanti, MN) that automatically detached the collar at the end of the study. Collar units recorded tri-axial acceleration data at 12 Hz during daylight hours (06-18h). Data used here include a single day from each of five adult females weighing between 14 and 16 kg.

Koala fieldwork was undertaken on private land at Cape Otway, Victoria, Australia. From 1 November 2010 to 24 January 2011, 27 adult koalas were captured using a standard 'noose and flag' method. Each koala was fitted with a VHF radio collar (Sirtrack, Havelock North, NZ; 100 g). A tri-axial accelerometer data logger (Gulf Coast Data Concepts LLC, Waveland, MS 39576; model X6-1A; 55 g) was taped to the collar so that the x-axis ran along the spine of the animal. Koalas were processed (without anesthesia) at their capture points and then released. Each koala was located at least twice (morning and evening) every day and then recaptured for collar removal after 7 days. Methods were approved by the Deakin University Animal Ethics Committee (A31-2010) and conducted under permit (10005379) by the Victorian Department of Sustainability and Environment.

Mountain brushtail possum fieldwork was undertaken in the Strathbogie Ranges, Victoria, Australia between spring 2011 and winter 2013. Possums were captured in large wire cage traps, placed on the ground, and baited with peanut butter and apple. Individuals were anaesthetized with an intramuscular injection of tiletamine/zolazepam (6 mg kg $^{-1}$; Zoletil ®; Virbac Australia, Peakhurst, NSW, Australia), fitted with a custom-built tracking collar incorporating a GPS (Mobile Action; model i-gotU GT-120), Accelerometer (Gulf Coast Data Concepts LLC, Waveland, MS 39576; model X8), and VHF transmitter (Sirtrack, Havelock North, NZ) with a total mass < 80 g and released at dusk at their capture site. Collars were removed by tracking individuals via VHF, re-capturing in wire cage traps, and removing collars without anesthesia before immediately releasing individuals at the capture site. A total of 32 individual possums weas tracked for 2-7 nights at a time, totaling 149 nights of data across the 8 consecutive seasons. Methods were approved by the Deakin University Animal Ethics Committee (A45-2011, B33-2012) and conducted under permit by the Victorian Department of Sustainability and Environment

Female Red deer were collared in Doupov and Slavkov Mts, Czech Republic within their winter enclosure sites between January 2019 and May 2019. Deer were baited to a central area within the enclosures and were anesthetized using a mixture of ketamine and xylasin delivered by dart injection Pneudart 3 mL. After darting, deer were tracked visually and using spoor. Once the individual was located, it was visually evaluated from a distance by a trained field biologist to determine if handling and collaring was safe for the deer and field crew. If so, the individual was blindfolded using a cloth bag to reduce stress while field biologists secured the animal’s limbs and neck to prevent injury. Deer were then fitted with a GPS wildlife collar (Vectronic

Aerospace GmbH, Germany) equipped with a built-in tri-axial accelerometer and magnetometer programmed to record continuously at 10 Hz (Wildbyte Technologies) and weighing 750 g (lifetime *ca.* 1 year). Deer were monitored until conscious by a field biologist who confirmed that collared individuals left the area under their own power without obvious impairment. Collars contained breakaway mechanisms allowing the data to be recovered and uploaded for analysis. Data used in the current analyses were collected after the deer had been released from their winter enclosures (1.5.2020). All handling, anesthesia and collaring procedures were approved by the ethics committee of the Ministry of the Environment (number MZP/2018/630/694).

Wild boar were collared in Kostelec nad Cernymi lesy, Czech Republic between May 2019 and January 2020. Large circular coral traps baited with corn were set each trap night and were equipped with a custom-designed camera system programmed to send images to researchers immediately after a boar had triggered the trap door. Selected adult female boar were then anesthetized using a mixture of ketamin, xylasin, tletamin and zolezapam delivered by dart injection (Pneudart 3 mL)). Once recumbent, the coral door was opened to release any non-anesthetized individuals and to allow us to fit anesthetized females with a GPS wildlife collar (Vectronic Aerospace, GmbH, Germany) equipped with a built-in tri-axial accelerometer and magnetometer sensor programmed to record continuously at 10 Hz (Wildbyte Technologies) (total weight 750 g; lifetime *ca.* 1 year)). The animals were monitored until conscious by a field biologist who confirmed that collared individuals returned to the forest under their own power without obvious impairment. Collars contained breakaway mechanisms allowing for the data to be recovered and uploaded for analysis. All handling, anesthesia and collaring procedures were approved by ethics committee of the Ministry of the Environment (number MZP/2019/630/361).

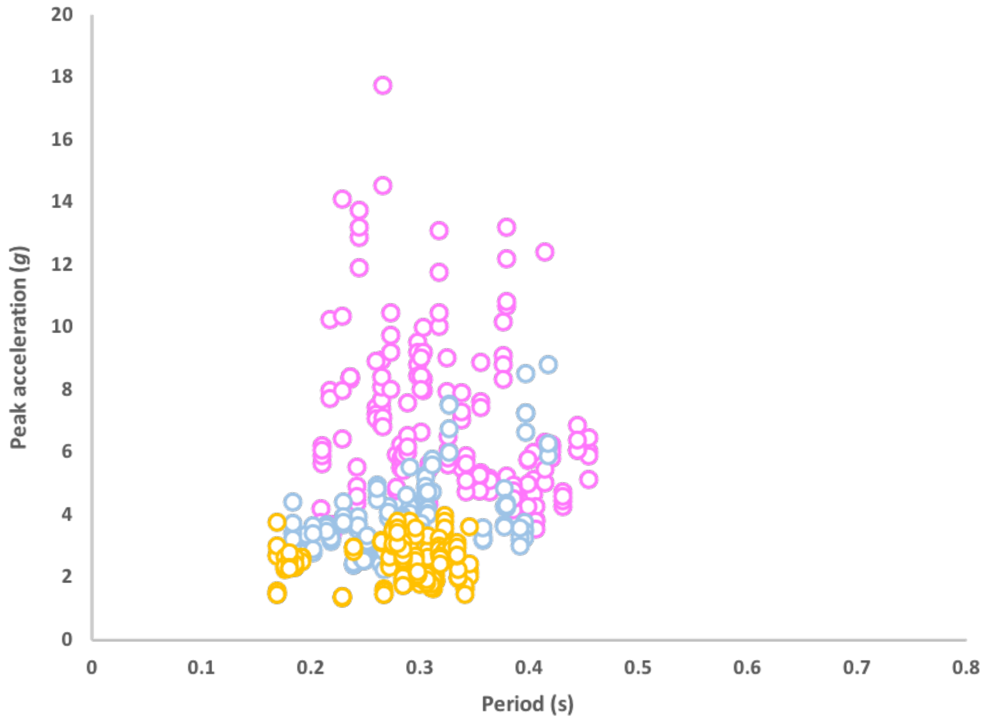


**Fig. S1.**

Peak amplitudes of (the vectorial sum of) accelerations versus period between peaks for walking (orange), trotting (blue) and bounding (pink) in twelve domestic dogs ranging 2-45 kg. Datapoints are taken from the four greatest peaks in acceleration per 20 m trial per dog.

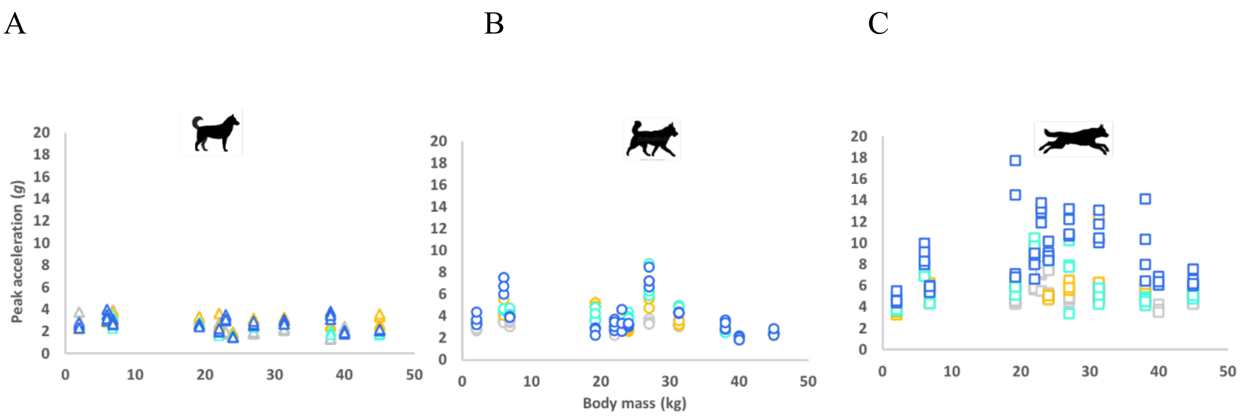


**Fig. S2.**

Peak amplitudes of (the vectorial sum of) accelerations versus body mass for twelve dogs ranging 2-45 kg in body mass. A) walking. B) trotting. C) bounding. Data points include the four greatest peaks in acceleration per 20 m trial per dog and are colored according to tag mass as a percentage of carrier body mass; 0% (grey), 1% (yellow), 2% (light blue) and 3% (dark blue).

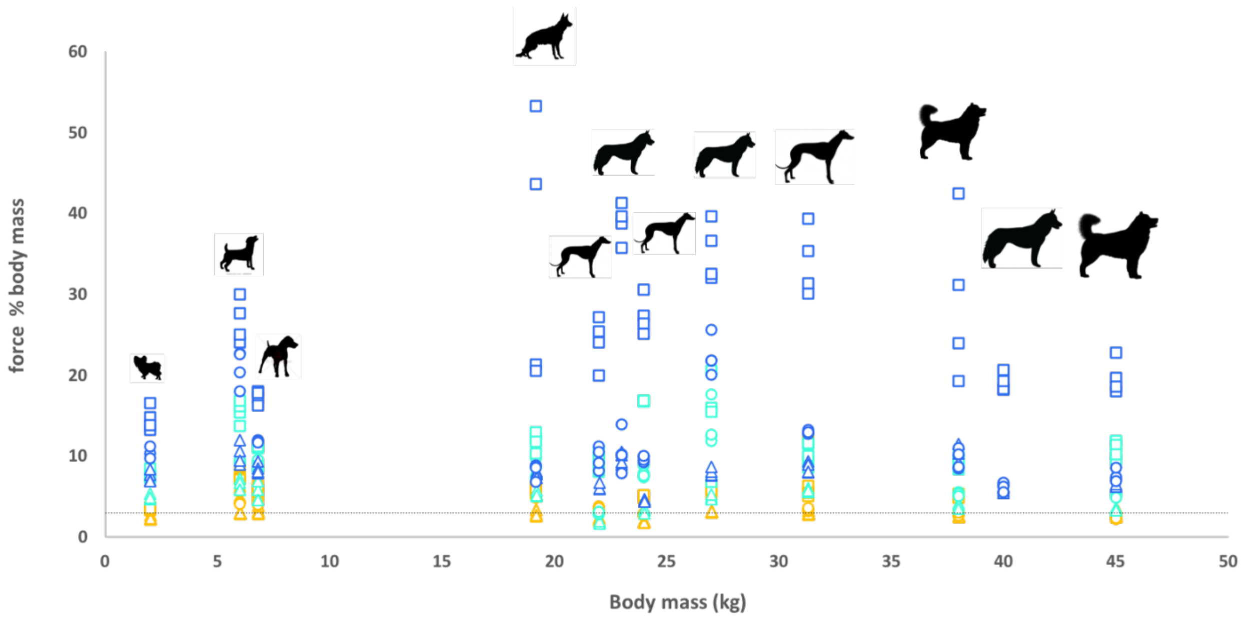


**Fig. S3.**

Forces exerted by the tags as a percentage of dog body mass for twelve dogs ranging 2-45 kg in body mass. Data points show walking (triangles), trotting (circles) and bounding (squares), and are colored according to tag mass as a percentage of carrier body mass; 1% (yellow), 2% (light blue) and 3% (dark blue). The dashed grey line indicates the relative force of a tag of 3% body mass at 1 *g*.

**Table S1.**

Percentage time the tags were exposed to accelerations (vector sum) greater than the values specified in brackets according to species and gait. Note that pine martens only moved substantially by bounding.

| Species | Walk (1 *g*) | Trot (1 *g*) | Trot (2 *g*) | Bound (1 *g*) | Bound (2 *g*) | Bound (3 *g*) |
|---|---|---|---|---|---|---|
| African lion | 31% | 46% | 12% | 62% | 33% | 14% |
| Cheetah | 31% | 60% | 19% | 88% | 60% | 33% |
| European badger | 31% | 47% | 9% | 44% | 27% | 14% |
| Pine marten | - | - | - | 42% | 23% | 11% |

**Table S2.**

Mean peak accelerations (SD) [vectorial sum – $g$] per stride measured using a collar-attached tag for four wild animal species as a function of activity. Note that pine martins only moved substantially by bounding

| Species | Walk | Trot | Bound |
|---|---|---|---|
| African lion | 1.81 (0.13) | 3.08 (0.37) | 4.73 (0.88) |
| Cheetah | 1.37 (0.05) | 2.70 (0.39) | 6.25 (0.79) |
| European badger | 1.39 (0.17) | 2.09 (0.34) | 3.86 (0.68) |
| Pine marten | - | - | 3.67 (0.52) |

**Table S3.**

Final outputs of linear mixed-effects models conducted to investigate the factors influencing peak accelerations in wild carnivores. Data were logged to ensure that model residuals were normally distributed.

| **Parameter** | **Final model terms** | ***F*** | **DF** | ***P*** | ***$R^2$ fixed*** | ***$R^2$total*** |
|---|---|---|---|---|---|---|
| log peak acceleration | log(period) | 0.013 | 1, 210.40 | 0.908 | 0.88 | 0.92 |
| | gait | 1083.07 | 2, 208.39 | <0.0001 | | |
| | body mass | 3.00 | 1, 18.79 | 0.100 | | |
| period | body mass | 99.98 | 1, 14.46 | <0.0001 | 0.73 | 0.76 |
| | gait | 114.50 | 2, 210.03 | <0.0001 | | |
| | bodymass x gait | 76.27 | 2, 209.29 | <0.0001 | | |

**Table S4.**

Summary of the outputs of linear mixed-effects models conducted to investigate how dog body mass ($M_b$), collar mass as a percentage of carrier mass ($\%M_b$), travel speed ($U$) and gait influence different parameters. Dog ID was included as a random effect in all models. Final models following the removal of non-significant interaction terms are reported

| Model | Parameter | Final model terms | *F* | DF | *P* | $R^2_f$ | $R^2_t$ |
|---|---|---|---|---|---|---|---|
| A | peak acceleration (*g*) | period | 10.81 | 1, 496.55 | 0.001 | 0.19 | 0.54 |
| | | gait | 340.88 | 2, 509.97 | <0.0001 | | |
| | | body mass | 0.21 | 1, 10.15 | 0.658 | | |
| | | period x gait | 7.92 | 514.49 | <0.0001 | | |
| B | period between peaks (s) | body mass | 3.37 | 1, 9.98 | 0.096 | 0.20 | 0.54 |
| | | gait | 47.54 | 2, 508.02 | <0.0001 | | |
| | | body mass x gait | 9.90 | 2, 508.04 | <0.0001 | | |
| C | speed (m $s^{-1}$) | body mass | 5.64 | 1, 9.96 | 0.039 | 0.82 | 0.86 |
| | | % body mass | 1.67 | 3, 507.49 | 0.172 | | |
| | | gait | 1500.614 | 2, 505.04 | <0.0001 | | |
| | | body mass x gait | 0.15 | 2, 505.10 | <0.0001 | | |
| D | peak acceleration (*g*) | speed | 40.59 | 1, 506.91 | <0.0001 | 0.66 | 0.74 |
| | | % body mass | 35.15 | 3, 499.60 | <0.0001 | | |
| | | gait | 21.15 | 2, 502.31 | <0.0001 | | |
| | | body mass | 3.51 | 1, 10.12 | 0.090 | | |
| | | speed x % body mass | 4.44 | 3, 500.77 | 0.004 | | |
| | | % body mass x gait | 4.33 | 6, 498.57 | 0.0002 | | |
| E | force as % body mass | body mass | 4.62 | 1, 9.98 | 0.057 | 0.38 | 0.84 |
| | | % body mass | 126.44 | 2, 356.97 | <0.0001 | | |
| | | gait | 75.28 | 2, 356.01 | <0.0001 | | |
| | | body mass x % body mass | 37.68 | 2, 356.90 | <0.0001 | | |
| | | body mass x gait | 7.38 | 2, 356.03 | <0.0001 | | |
| | | % body mass x gait | 16.10 | 4, 356.01 | <0.0001 | | |

**Table S5.**

Dog ID and morphometrics

| Name | Breed | Sex | Mass (kg) | Body (cm) | Fore (cm) | Hind (cm) |
|---|---|---|---|---|---|---|
| Jess | Papillon | F | 2.0 | 25 | 30 | 20 |
| Daisy | Jack Russel | F | 6.0 | 35 | 29 | 29 |
| Faith | Patterdale | F | 6.8 | 36 | 28 | 30 |
| Rags | Alsatian x Staffordshire Bull Terrier | M | 19.2 | 49 | 48 | 50 |
| Pip | Lurcher | F | 22.0 | 62 | 55 | 54 |
| Luna | Husky | F | 23.0 | 53 | 52 | 50 |
| Barnaby | Lurcher | M | 24.0 | 57 | 60 | 61 |
| Snow | Husky x Malamute | F | 27.0 | 62 | 59 | 59 |
| Saffy | Lurcher | F | 31.3 | 63 | 48 | 55 |
| Maya | Husky x Malamute | F | 38.0 | 67 | 62 | 61 |
| Molly | Husky | F | 40.0 | 60 | 59 | 57 |
| Malcom | Husky x Malamute | M | 45.0 | 73 | 63 | 59 |
